# Ancient DNA from chewing gums connects material culture and genetics of Mesolithic hunter-gatherers in Scandinavia

**DOI:** 10.1101/485045

**Authors:** Natalia Kashuba, Emrah Kirdök, Hege Damlien, Mikael A. Manninen, Bengt Nordqvist, Per Persson, Anders Götherstörm

## Abstract

The discussion of an early postglacial dual-route colonization of the Scandinavian Peninsula is largely based on associating genomic data to an early dispersal of lithic technology from the East European Plain. However, a direct link between the two has been lacking. We tackle this problem by analysing human DNA from birch bark pitch mastics, “chewing gums”, from Huseby Klev, a site in western Sweden with eastern lithic technology. We generate genome- wide data for three individuals, and show their affinity to the Scandinavian hunter-gatherers, or more precisely, to individuals from postglacial Sweden. Our samples date to 9880-9540 calBP, expanding the temporal range of this genetic group as well as its distribution. Human DNA from mastics provides a clear connection between material culture and genetic data. We also propose that DNA from different types of mastics can be used to study environment, ecology, and oral microbiome of prehistoric populations.

---

Writing human history using lines of evidence from both ancient DNA and archaeology, has been debated since the first DNA-sequence was recovered from human remains ^1–3^. Central to this discussion is that ancient human skeletal material often comes from ritual contexts, many times without associated artefacts, and therefore lacks any clear connection to the material that archaeologists use to study the life of past societies. Here we suggest that a solution to this problem lies in obtaining human aDNA directly from the remnants of material culture, more specifically from “chewing gums”, masticated lumps, often with imprints of teeth and/or fingers ^4^ (Fig. 1A). The material we use is made of birch bark pitch, which is known to have been used as an adhesive substance in lithic tool technology and as a cement to seal wood and pottery vessels in prehistoric Eurasia (Supplementary Note 1).

**Fig. 1.**
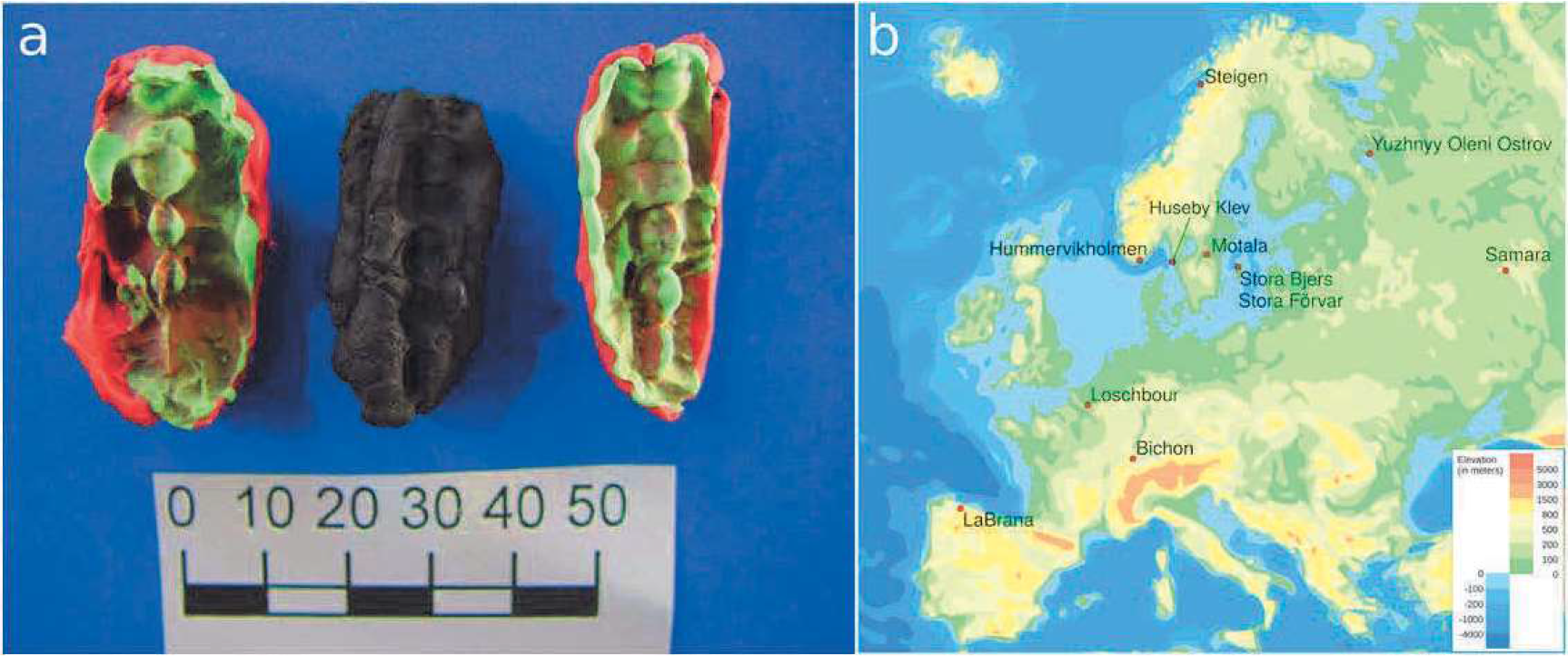
The studied material and its origin, **a** One of the chewing gums from Huseby Klev, with two plastelina casts showing positives of teeth imprints. One cast for each side of the ancient mastic piece. The cast to the left is of the backside, the one to the right is of the side facing up on this photo. Scale bar: 50 mm (photo by Verner Alexandersen). **b** The location of Huseby Klev and the places from which the other ancient genomes used in the demographic history analyses originate.

In this study, we investigate the relationship between Stone Age populations and cultural traits expressed in lithic technology in Scandinavia. Earlier aDNA studies suggest the presence of three genetic groups in early postglacial Europe: Western hunter-gatherers (WHG), Eastern hunter-gatherers (EHG) and Scandinavian hunter-gatherers (SHG)^5^. The SHG have been modelled as a mixture of WHG and EHG ^5–7^. SHG is genetically the most diverse, suggested to be a consequence of an immigration which took place around 10,300 calBP ^7^. This is consistent with the rich archaeological evidence of a dual-route human dispersal into the Scandinavian Peninsula at the end of the latest Ice Age: first migration from the south, which took place at c. 11,500 calBP, and a second one from the northeast, detected at about 10,300 calBP ^8–11^. These migrations are associated with differing lithic technological traditions, the one from the later migration being connected to the pressure blade technology, known in preceding centuries from the East European Plain, Karelia, and Finland, and suggested to have rapidly spread into Scandinavia along a north-eastern route. The connection between a migration from the north-east into Scandinavia and the significant technological change in the form of pressure blade technology, remains a suggestion, as none of the SHG individuals studied for aDNA have been directly linked to the early eastern blade production technology.

We explore the connection between the demography and material culture of Scandinavian hunter-gatherers by studying mastics and remains of lithic tool production from the Mesolithic Huseby Klev site (Fig.1 B). At this location in western Sweden, chewing gums and other pieces of mastic, along with lithic remains, are found in a temporally well defined context, the “deep pit” excavation trench, dated to c. 10,040-9610 calBP (Supplementary Note 1). Analysis of the lithic material shows that the eastern tool technology was used already at an early phase of the site. We consequently generated genome-wide data from mastics, representing three individuals and use the results to reevaluate the earlier suggested co-dispersal of humans and eastern lithic technology.

## Results

### Extracting and authentication of ancient DNA from ancient mastics

We tested two different aDNA extraction methods on eight mastic samples in total. First we performed Yang-Urea extraction following the modifications employed by Svensson et al. ^12^ of the published protocol^13^. We named the successful samples ble004, ble007 and ble008. For the ble008 sample we also used an extraction kit designed to process samples with high inhibitor content (QIAamp PowerFecal DNA Kit, Qiagen), with slight modifications to the protocol provided with the kit. Blunt-end libraries were built on the concentrated DNA ^14^ which were amplified and shotgun sequenced at the Science for Life laboratories (SciLife) in Stockholm. Initial tests showed that the individual libraries from Yang-Urea extracts contained authentic ancient human DNA, ranging from 2% to 8%. The library built on the PowerFecal DNA Kit extract contained 23% endogenous DNA. The 20-fold increase in endogenous DNA content makes the PowerFecal DNA extraction kit a valid approach to process mastic samples. By repeating the process with successful extracts, we obtained genome-wide data from three of the mastic pieces, ranging from O.lx to 0.49x coverage (Table 1). We merged individual libraries using *SAMtools* ^15^ and calculated library statistics (before and after PMD filtering, Supplementary Table 1, 2) and produced damage plots (Supplementary Figure 1).

**Table 1:** The library properties for *ble* samples after processing and pmd filtering.

| Sample | ble004 (merged) | ble007 | ble008 (merged) |
| --- | --- | --- | --- |
| Human Sequences | 6,463,893 | 5,802,458 | 18,393,636 |
| Read length (mean) | 66.95 | 76.98 | 95.57 |
| Nuclear genome coverage | 0.11 | 0.10 | 0.49 |
| MT contamination estimate | 3.08 | 1.23 | 8.61 |
| MT haplogroup | U5a2d | U5a2d | U5a2d |
| Xseq | 261,823 | 170,651 | 866,414 |
| Yseq | 22,093 | 69,025 | 26,051 |
| Biological Sex | XX | XY | XX |
| Number of SNP (compared with Human Origins dataset) | 20,103 | 19,665 | 52,211 |

### Contamination and pmd filtering

We estimated mitochondrial contamination rates using near-private consensus alleles as described by Green et al. ^16^ To exclude the effects of sequencing errors, we used bases that have a base-call and mapping quality score of more than Q30. Also, we filtered the positions where we detected transition patterns to compensate the post-mortem damage (Supplementary Table 3).

We used the *PMDtools* software to filter out the possible contaminant sequences ^17^. This tool compares each aDNA sequence to its modern counterpart to calculate deamination specific nucleotide transitions and assigns a *pmd score*. This score is used to evaluate the authenticity of the sequence. We set the *pmd* threshold to 0 and removed contaminant DNA sequences below this threshold (Supplementary Figure 2). After removing the potential contaminants from the merged libraries, we re-calculated the library statistics, deamination patterns, and MT contamination estimates to analyze the authenticity of the dataset. Deamination patterns and MT contamination rates (Table 1) present a strong case that the aligned data is authentic and represents the individuals that chewed the ancient mastic (Supplementary Table 2, 4).

### Relationships between ancient individuals

We used READ ^18^, to explore kinship between the individuals. READ compares the non-overlapping 1Mb segments in the genome and calculates the non-identical allele ratio between the samples (PO). Lower PO values mean more shared chromosomal segments. We confirmed that none of the genomes are identical to each other (Supplementary Table 5). We also found that ble004 and ble007 have a possible second degree relationship. However, it should be noted that using three individuals for analysis is not recommended for this tool, and we refer form using this result in further discussion. In summary, we can confirm that we sequenced DNA from three distinct individuals.

### Mitochondrial DNA

We used *samtools mpileup* command to create mitochondrial consensus sequences using the nucleotides that have a base-call and mapping quality score of more than Q30. We assigned mitochondrial haplogroups with HaploFind (a) and HaploGrep 2 (b) (Table 1). We reviewed the results with PhyloTree (build 17) (c). Mitochondrial genomes from all three individuals belong to the U5a2d haplogroup. ble008 was assigned to U5a2 by HaploGrep 2, but the same sequence is assigned to U5a2d haplogroup by HaploFind and we accept this result (Supplementary Excel Table 1). The mitochondrial U5a2d haplogroup is consistent with earlier published results for ancient individuals from Scandinavia, U5a being the most common within SHG. Of the 16 Mesolithic individuals from Scandinavia published prior to our study, seven belong to the U5a haplogroup, nine share the U2 and U4 haplogroups^6,7^.

### Demographic history and population genomics of Huseby Klev individuals

To explore the demographic history of the ancient individuals, we curated a set of ancient genomes (Supplementary Table 6) and compared with the publicly available Human Origins SNP reference set^19,20^. This set contains 594,924 SNP’s from 2,404 modern individuals and 203 different worldwide populations. We coded deamination patterns as missing data to compensate for possible biases caused by deamination patterns.

We used principal component analysis (PCA) to acquire an overview of the affinity of the ble individuals with selected ancient and modern populations ^21^. We merged ancient individuals with the Human Origins reference set, coded nucleotide transitions as missing data, and used Procrustes transformation to project ancient individuals on the principal component space (Supplementary Figure 3). The projected ancient individuals show close affinity to modern day North, East, and Western European populations, and *ble* individuals from Huseby Klev (the earliest among the SHGs) cluster with the ancient genomes originating from Scandinavia ^5^. These three samples are located between the two Hummervikholmen individuals (Norway), and the Stora Forvar SF9/12 (Sweden), all three with dates earlier than c. 9000 calBP. By reproducing EHG and WHG populations in this plot^7^, we confirm the close affinity of ancient individuals from Scandinavia to WHG and EHG (Fig 2a).

**Fig. 2.**
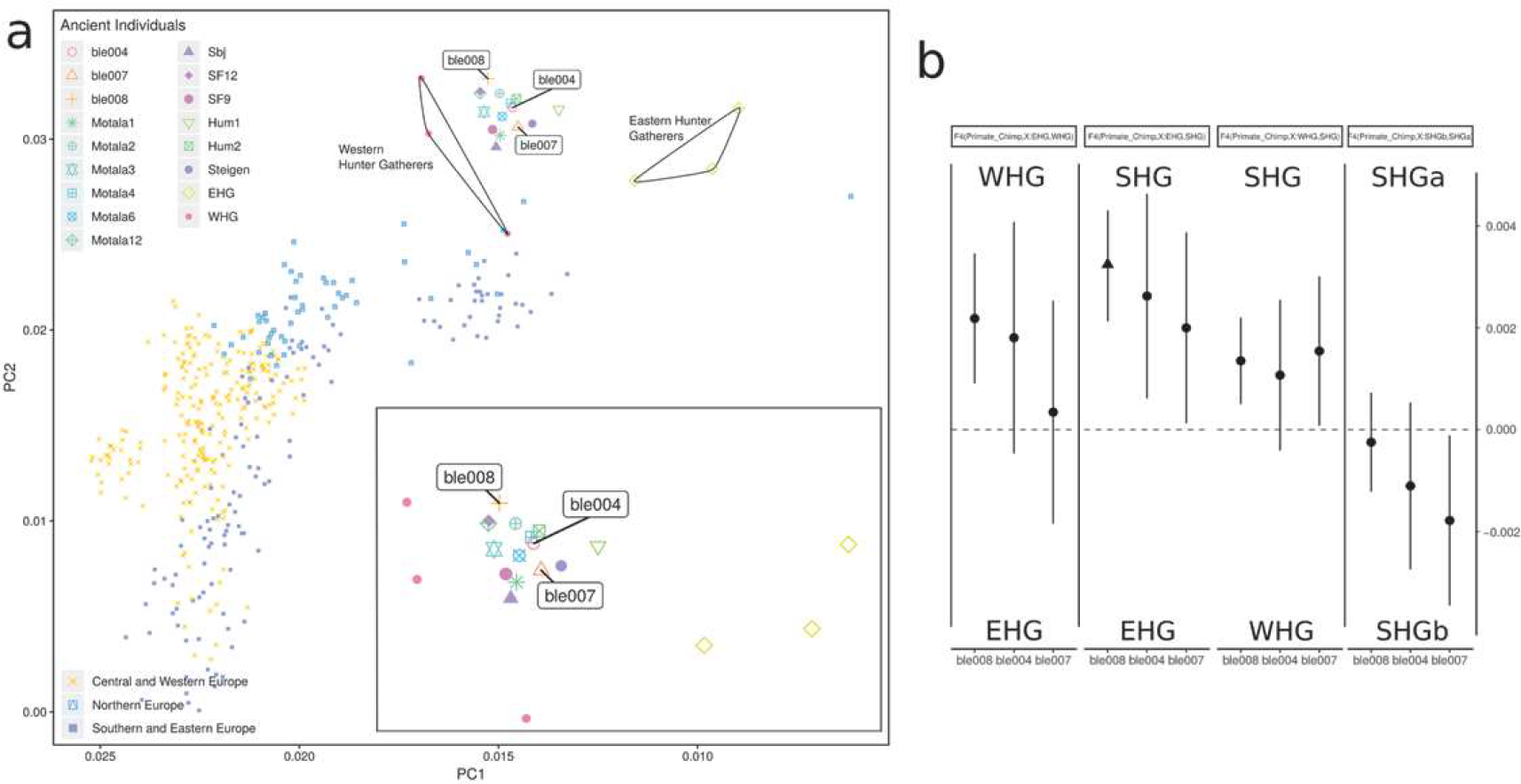
Principal component analysis of the Huseby Klev individuals within the diversity of Mesolithic individuals from Europe, **a** The magnified section incaptures BLE individuals’ relation to WHG, EHG and SHG individuals (Supplementary Table 6). **b** Results of relative allele sharing (F4) test between Huseby Klev individuals and ancient population groups (the triangle shows the significant deviations from zero)

To evaluate the relationship between the Huseby Klev individuals and other Mesolithic Scandinavians, we examined the relative shared allele frequencies and estimated shared drift among populations via performing D and F4 tests. These tests propose formal statistical frameworks to study the patterns of allele frequency correlation across populations ^20^. We first tested the relative allele sharing of ble individuals between EHG and WHG groups. Results show a high contribution of WHG ancestry to BLE individuals (The highest contribution observed in ble008 individual F4: 0.022, Z-Score 1.8, Fig 2A). We compared the contribution between ancestry of EHG to SHG and WHG to SHG for BLE individuals. Results show that all tested individuals have relative allele sharing with the SHG group, with ble008 individual showing the significant value (F4: 0.03, Z-score:2.95). We divided the SHG group into two groups: SHGa and SHGb (ancient individuals found in contemporary Norway and Sweden, respectively). We based this on both the geographical distribution and the previous studies demonstrating the close relation of SHGa to EHG group and SHGb to WHG group ^7^. To further explore the demography within the SHG group, we compared the ancestry of BLE individuals within SHGa and SHGb groups. This comparison revealed a high relative shared drift between BLE individuals and the SHGb group (Supplementary Excel Table 2, for D and F4 tests).

To support our findings from the D and F4 tests, we used the model-based clustering algorithm admixture ^22^, which is helpful when exploring the genetic components in a given dataset. The essence of this algorithm is to calculate the admixture proportions as a parameter of a model. The results show that starting from K15, there is a similarity in genetic elements between the BLE individuals and WHG, and SHGb populations (Fig. 3, Supplementary Figure PDF 1).

**Fig. 3.**
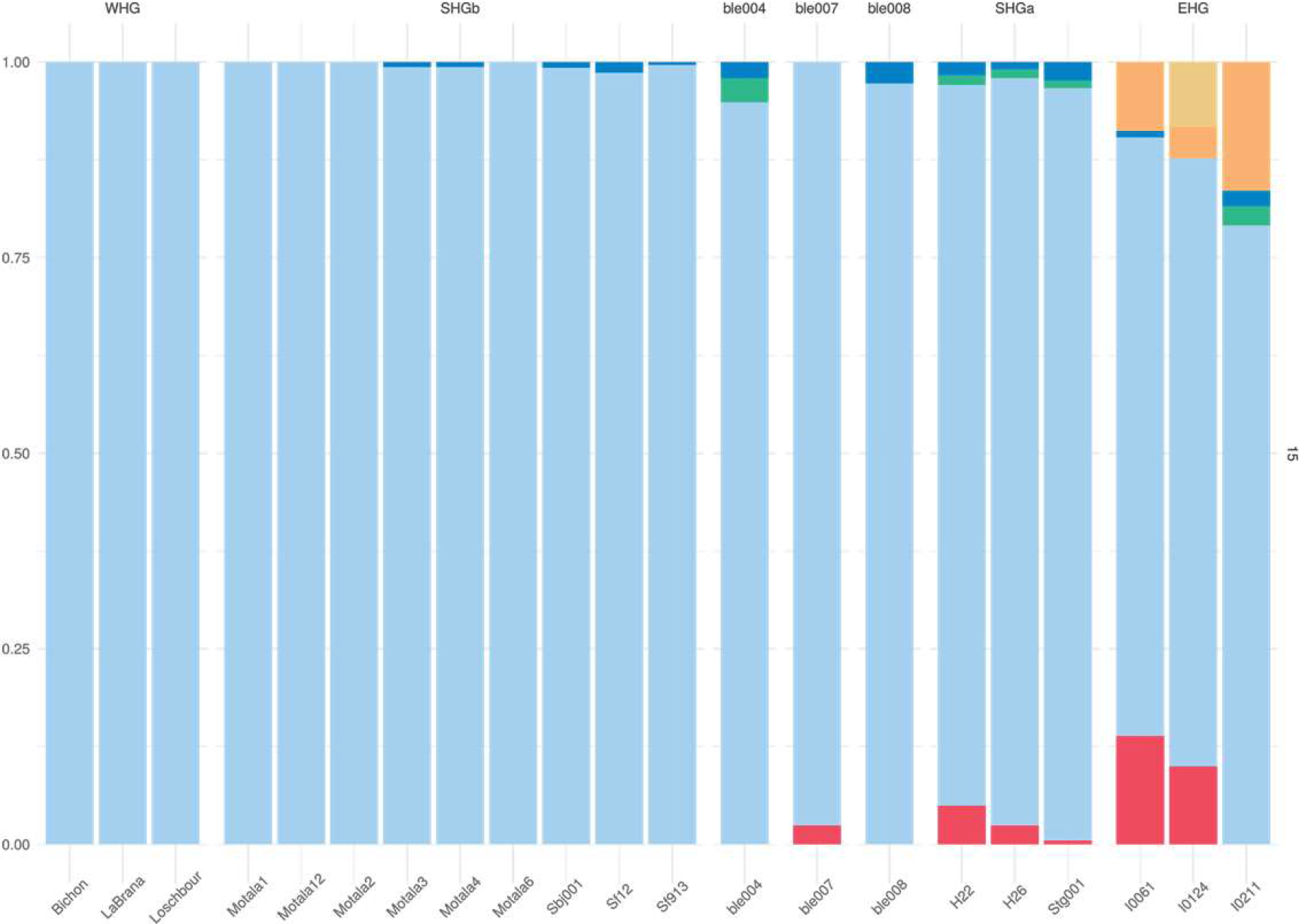
Admixture analysis showing the major mode for K=15. The figure represents 11 runs out of 20 replicates (Greedy algorithm ran with the Jaccard distance and a 0.97 similarity threshold)

### Lithics results

The technological analysis conducted for the purpose of this study, shows that the lithic artifacts from the deep pit display clear affiliation with the eastern pressure blade technology, as documented from a large number of sites in northern and western Scandinavia, eastern Fennoscandia, and the East European Plain ^9–11, 23^. No artefacts diagnostic to the preceding Early Mesolithic blade technology, that would indicate chronological or technological mixing, were observed (Supplementary note 1).

Based on the composition of lithic artefact types, the site appears to represent a production site to which lithic raw materials, in their more or less unworked condition, were transported, and where initial core preparation and exploitation was performed. Additionally, some standardized blade production and re-tooling was performed on-site, visible in the presence of discarded tools (a relatively low number) and regular blade blanks.

Blades were produced by the same overall concept: serial production from single-platform, sub-conical and conical cores with faceted and smooth platforms. No complete regular blade cores are present, but fragments of conical cores with visible scars deriving from the detachment of very regular thin blades, along with core rejuvenation flakes with small-flake faceting and a platform to front angle close to 90°, suggest that the eastern pressure blade technology concept was employed. Although the majority of the studied blades display features found in blades produced by direct and indirect percussion techniques, a selection of blades display diagnostic characteristics of the pressure technique, and the variation in knapping techniques is best explained as related to the different stages of the production process (Supplementary note 1). Morphometric analysis shows the production of a consistent range of blade blanks, which in turn allowed the production of standardized tools, such as barbed points (hulling-type), slender lanceolate microliths, as well as blades with lateral retouch on one or both edges. The last mentioned were probably used as inserts in composite slotted tools, to which the inserts were attached using mastic made of birch bark pitch ^24^ (Fig. 4). A bone point with remains of pitch retrieved from the deep pit shows that birch bark pitch mastic was part of tool production at the site, while fragments of slotted points, contextually dated to the same period as the finds from the deep pit, were found nearby^25^.

**Fig. 4.**
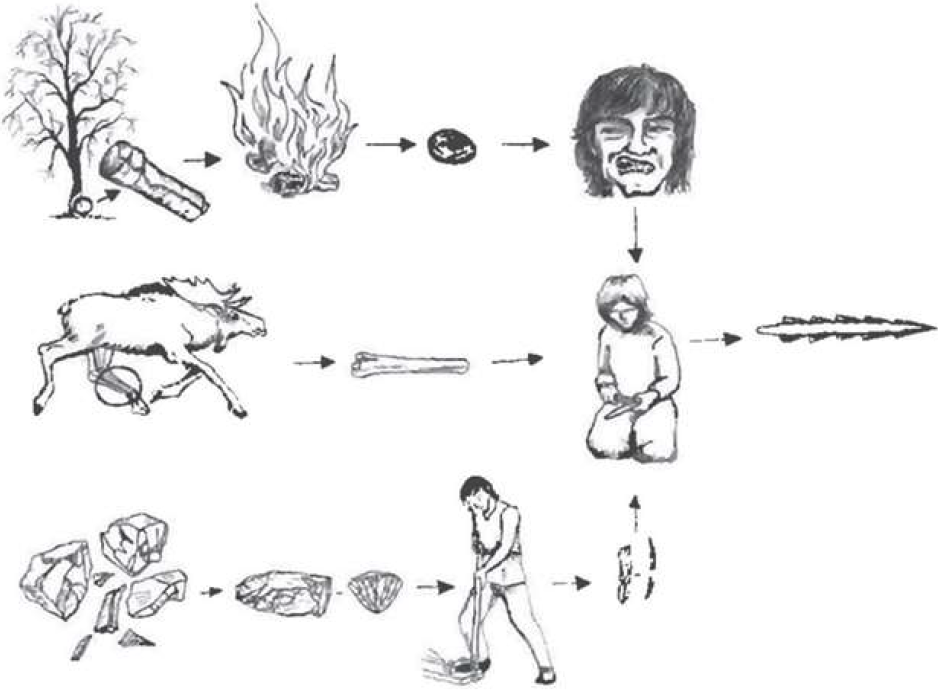
Operational chains used in the processing of raw materials during composite bone point production. Lithic blades served as inserts and birch bark pitch was used as an adhesive agent. Drawing: Kristina Steen ^26^ (with permission from Universitetsforlaget Oslo)

## Discussion

Prior to our study, ancient DNA has been retrieved from biological remains of ancient individuals: bones, mummified tissue^27^, hair^28,29^, and coprolites^30^. Human aDNA in soil samples has also been discovered and processed, but so far only to determine the presence of homo species^31,32^. Our results from ancient mastic add to the available sources for genomic data on ancient individuals. DNA from saliva, preserved in the mastics, yields ancient human aDNA, the authenticity of which we can confirm by studying damage patterns, contamination rates and population genomic analysis (Supplementary Figure 1, Supplementary Table 4, Supplementary Figure 3).

The genomic data from the mastics allows us to determine a close affinity between individuals from Huseby Klev and the previously defined SHG genetic group (Fig 2.). F4- and D-statistics confirm a close affinity to SHG, or more precisely, individuals found in Sweden (the Mesolithic SHGb subgroup). Using admixture analysis, we show that the studied Huseby Klev individuals have more of the WHG than the EGH component (Fig 2.), which is consistent with the genomic composition of the SHGb individuals. The Huseby Klev individuals allow us to root the SHGb group and extend its chronological span, as well as geographical distribution.

Prior to this study, genetically defined SHG individuals either had no association with lithic technology (SHGa: Steigen and Hummervikholmen) or the associated artifacts were not sufficient to make inferences about stone tool production technology known to the buried individual (SHGb: Stora Bjers). The only two links established between genetics and stone tool technology are found in the Stora Forvar cave’s Mesolithic layer (SHGb) and at the Motala Kanaljorden site (SHGb). The Motala Kanaljorden lithic inventory has characteristics typical of the handle core technology ^33^, a blade production technology common in Late Mesolithic Scandinavia, and different from the eastern blade production technology^9,32^. In Stora Förvar, the lithic inventory has techno-typological characteristics of a technology typical to the first postglacial pioneers entering Scandinavia from the south, including blade production from single- and dual platform cores by direct percussion techniques ^34^. Table 2 shows the different lithic technological traditions, all found within the genetically defined SHGb group (as a context with SHGa and stone tools is currently lacking). The earliest documented technology (context dated to 10,040-9610 calBP), associated with SHG, is the eastern blade technology visible in the production waste from Huseby Klev. This variety of technological traditions within the SHG group reminds us that genetic and cultural features are compatible only to some degree, and that we should be careful when merging information on cultural evolution and demographic processes.

**Table 2.** Scandinavian Mesolithic sites where both human aDNA and lithic artefacts are found. “Context date” is based on calibrated radiocarbon dates. Individual dates and the principle used in determining a context date are given in Supplementary note 2. “Genetic group” refers to a subdivision of the SHG group into SHGa and SHGb.

| Site | Context date | Genetic group | Blade production technology |
| --- | --- | --- | --- |
| Huseby Klev | 10,040-9610 calBP | SHGb | Eastern pressure blade technology |
| Stora Förvar | 9170-8000 calBP | SHGb | Maglemosian technological group 1&2 |
| Motala, Kanaljorden | 7760-7520 calBP | SHGb | Handle core technology |

The results from Huseby Klev allow us to finally connect the SHG group with the eastern pressure blade technology. However, the higher genetic affinity between Huseby Klev individuals and the WHG group challenges the earlier suggested tie between eastern technology and EHG genetics. Our results suggest either early cultural transmission, or a more complex course of events involving both non- and co-dependent cultural and genetic admixture.

By combining genomic data and the archaeological dwelling site context, we are able to gain new insights into the Mesolithic society and discuss the social organisation of of past populations. The fact that each of the studied mastic pieces was chewed only by single individuals, both male and female, and that the mastication of birch bark pitch was most likely connected to the process of tool-making and maintenance (an interpretation supported by the evidence of core processing, re-tooling, and hunting found at Huseby-Klev), allows for a discussion of gender roles within Mesolithic society. Combined with the fact that several mastics have imprints of deciduous teeth (Alexandersen 2005), the new information allows us to discuss gender in childhood. The possible interpretations are that tool-production was not restricted to one sex, or, if the individuals examined were children, that young individuals were not treated as males or females. When results from other dwelling sites besides Huseby Klev start to accumulate, we will be able to discuss the social organisation of past populations on a wider scale.

It is known that birch bark pitch mastics, but also mastics of other materials, have been widely used around the world from the Middle Palaeolithic onwards (Supplementary note 1), including in regions where human remains are not available to study, either due to bad preservation (e.g., large parts of Fennoscandia), or restrictions for the use of human remains (such as the “Kennewick Man conflict”^35^). In these situations mastic pieces offer a possible source for DNA. In addition, mastics are expected to be a source of information concerning the environment, ecology, and oral microbiome of prehistoric populations.

## Methods

### Mastics of birch bark pitch in Huseby Klev

A number of the Huseby Klev mastics bear traces of human teeth, while all of them have a chewing-gum like morphology, a dark colour, and a glossy surface. While modern experiments show that relatively simple methods can be used in the production of birch bark pitch (Supplementary note 1), the production technology is somewhat knowledge-intensive. It is mostly for this reason that teeth marks in the pitch are often considered as indicative of processing and use, i.e., making the pitch more viscous and pliable, rather than a sign of purely recreational use as a chewing gum. It is known that birch bark pitch was used in hafting stone tools, and for attaching flint blades to slots in composite tools.

Of the 115 finds of pitch from Huseby Klev (Supplementary note 1), eight lumps from the Huseby Klev deep pit have been subjected to chemical analysis^36^. Seven of them turned out to be birch bark pitch while one did not give results. Alexandersen ^4^ has studied 10 lumps with tooth impressions and, by comparing to modern parallels of tooth development and wear, determined the age of the chewers in these cases to have been between 5-18 years. In addition, a piece with teeth impressions from both an adult and a child has been reported^37^. Other pieces of pitch from the deep pit show a variety of wood and cordage impressions ^38^.

### Sample preparation

We chose eight birch bark pitch mastic pieces for analysis. After the first screening, we continued working with three of the samples: *ble004, ble007, and ble008*. The samples were processed in the clean room facilities of the Archaeological Research Laboratory (AFL, Stockholm University), dedicated to ancient DNA work. The mastic pieces were irradiated in a crosslinker, at about 6 J/cm^2^ at 254 nm. The outer shell of the mastics was discarded to avoid surface contaminants. The powder for extraction was produced using a Dremel drill or a scalpel and collected into 2 ml tubes. The weight of the obtained powder varied between 66 and 191 mg.

### Extraction

To extract the DNA, we performed several incubations. At each incubation the samples were kept in rotation. First, the samples were pre-digested at 45°C for 15 minutes in 1000 uL of extraction buffer, consisting of Urea, EDTA (0.5M) (VWR) and 10 uL of Proteinase K (10mg/ml)(VWR) ^13^. A negative control was added during this step and taken through the work process. The supernatant from the predigestion step was removed and a fresh extraction buffer (same as above) with proteinase were added to the samples and left for digestion overnight at 37°C. The supernatant from this step was stored, and more extraction buffer and proteinase were added to the samples (same as above) and left rotating at 55°C for 4 H. The final supernatant was collected and combined with the previous one (around 2000 uL) and spun down to 100 uL using membrane filters (Amicon Ultra-4 Centrifugal Filter Unit with Ultracel-30 from Millipore). The extract was purified using MinElute spin columns and a buffer set (both Qiagen). We modified the Qiagen protocol (reducing the PE buffer volume to 600 uL and performing two elutions using 55 uL of the EB buffer) and obtained about 110 uL of extract. For ble004 sample two extracts were made from two different samplings.

We performed an extraction on ble008 sample using QIAamp PowerFecal DNA Kit (Qiagen), which is used to remove inhibitors in stool, gut and biosolid samples. We used 0.25 g of the sample and collected it directly into a Dry Bead Tube, containing garnet beads, which during vortexing mechanically disrupts cell walls. We added a negative control at the first extraction step. After adding the reagents according to the protocol we used a thermoshaker and vortexed the samples at 65°C for 30 minutes at the highest speed available. All buffers were added according to the protocol provided with the kit, besides the step 14, where we reduced the amount of C4 solution to 1000 uL. As the extraction was finished, the DNA got captured on a silica membrane of a spin column and purified, resulting in 100 uL of product.

### Libraries and sequencing

We built double-stranded blunt-end libraries, using the modified protocol by Meyer and Kircher ^15^. We used 20 uL of the extract and 30 uL for the USER enzyme pre-treated libraries. The master mix for the blunt end repair step contains 4 uL of Buffer Tango 10X with BSA (Thermo Scientific), 0.16 uL 25 mM dNTP mix (Thermo Scientific), 0.4 uL ATP 100 mM 25 umol (Thermo Scientific), 12.64 uL ddH20, 2 ul T4 Polynucleotide Kinase (10 U / uL) (Thermo Scientific) and 0.8 T4 DNA Polymerase (5 U / uL) (Thermo Scientific), and incubation step at 25°C for 15 min, followed by 5 min at 12°C. We used MinElute spin columns to purify the product, reducing the volume of the washing buffer PE to 600 uL. The ligation master mix contains 10 uL ddH20, 4 uL 10 × T4 DNA ligase buffer (Thermo Scientific), 4 uL PEG4000 50% (w/v) (Thermo Scientific), 1 uL of Adapter mix P5/P7 lOOmM (10 pmol) (Biomers.net) and 1 uL of T4 DNA ligase 5 weiss U /uL (Thermo Scientific), and is incubated at 22°C for 30 min. We purified the product as above and continued to fill- in the adapters with the following master mix: 14.1 uL of ddH20, 4 uL of 10 X ThermoPol reaction Buffer (BioLabs), 0.4 uL of 25 mM dNTP mix (Thermo Scientific) and 1.5 uL Bst DNA Polymerase Large Fragments 8,000 U/ml (BioLabs). The incubation steps are 37°C for 20 min, followed by 20 min at 80°C.

We performed UDG treatment before blunt end repair for several libraries ^39^. We used USER Enzyme 1,000 u/ml (BioLabs) and incubated the extract with the ingredients for a blunt end mastermix (excluding polymerase, which was added after the incubation) for 3 H at 37°C. We then proceeded with the blunt end protocol. For sample ble004 9 libraries were built, 5 of which have been pretreated by USER enzyme. For ble007 sample 5 libraries were produced, for ble008 sample 6 libraries.

Libraries were amplified with 10 μM index primers (Biomers.net), using AmpliTaq Gold 1000 Units 5 U / ul (Applied Biosystems) for blunt-end libraries, and AccuPrime™ Pfx DNA Polymerase (2,5U/ul) (Invitrogen) polymerase for damage repair libraries. We determined the number of cycles using quantitative PCR(qPCR), with reagents from Thermo Scientific and Biomers. Libraries were sequenced on the lllumina Hiseq X platform at the SciLife center in Stockholm.

### aDNA investigation and data processing

We merged paired-end reads if they had at least 11 nucleotides of overlap using the *MergeReadsFastCL_cc.py* script ^40^ and processed reads for any remaining adapter sequences. Afterwards, we treated each sequence as single-end read and aligned to human reference with custom parameters (*bwa aln* command with seeds disabled −I 16500 −n 0.01 −o 2) to allow more mismatches and gap events ^17,19,40^. We merged mapped libraries (*ble004_dr, ble004_nondr, ble007, ble008 and ble008 new method*) using *samtools merge*, and filtered reads that are PCR duplicates with *FilterUniqSAMConscc.py* script, having less than 90% sequence identity with the reference chromosome, smaller than 35 nucleotides, and mapping quality less than 30 ^40^. For *ble004*, we merged filtered damage repaired and non-damage repaired libraries to produce the final bam file.

### Principal component analysis

For this analysis, we merged the libraries of ancient individuals with Human Origins dataset separately by coding the nucleotide transitions as missing data. We used the *smartpca* program to calculate eigenvalues for each ancient individual. Then we used Procrustes transformation to project ancient individuals on the principal component space. Human origins reference genome set contains 594,924 SNP’s from 2,404 modern individuals from 203 populations worldwide ^19,20^. To compensate for the biases that could be introduced from PMD decay, we coded deamination transitions as missing data. PC plot with the entire Human Origins database is shown in Supplementary Figure 3.

### D and F4 statistics

To test the population affinities between the ancient individuals, we used the *popstats* program to calculate D and F4 statistics ^41^. These tests propose formal statistical frameworks to study the patterns of allele frequency correlation across populations ^20^. D tests provide evidence of significant deviations from a tree-like population structure, which could be demonstrated as ((A, B) (X, Y)). Positive D values indicate a population affinity between A, X and B, Y. Moreover, F4 tests give information about the direction of the shared genetic drift. Similarly, positive values indicate a shared genetic drift between A and X and B and Y. In both cases, negative values indicate a relationship between A, Y, and B, X. We followed the workflow as described in Skoglund et al. ^41^: We computed standard errors using a block jackknife weighted by the number of SNPs in each 5 cM and we reported Z-scores as normalized Z = D/s.e. Z>2 was interpreted as a significant deviation from zero. To calculate F4 statistics, we used the flag --f4 as described in the manual.

### Model-based clustering

We used a model-based clustering algorithm called *admixture* to understand the population structure in our dataset ^22^. To use our merged data with this tool, we first pseudo-haploidized the dataset by removing one allele randomly from the reference panel. Then we filtered our dataset for linkage disequilibrium using Plink with the parameters --*indep-pairwise 50 5 0.5* ^42^. We ran K=2 to K=20 with 20 different replicates using different random seeds. To detect common signals observed in independent admixture runs, we used a greedy algorithm implemented in *pong* software ^43^ (Supplementary figure PDF 1).

### Lithic technology

The lithic blade production concept at Huseby Klev was reconstructed by defining the production methods and knapping techniques used at the site. A dynamic-technological classification including a simplified *chaîme opératorie* analysis of the complete lithic assemblage and an attribute classification of a selection of the artefacts was employed as the methodological basis ^44–46^. In all 1849 flint artefacts from the deep pit were studied. Altogether 86 artefacts were considered high-priority in determining blade production methods and knapping techniques, and were subsequently selected and catalogued according to the attribute classification (Supplementary note 1).

### Data availability

The sequences are available at …

## Supporting information

## Acknowledgements

We express gratitude to Arielle Munters, who processed the raw DNA sequence data, and Vendela Kempe Lagerholm for support with the laboratory experiments. We also wish to thank the National Historical Museums in Sweden, Arkeologiska uppdragsverksamheten in Mölndal, represented by Glenn Johansson providing practical assistance during the lithic analysis. Emrah Kirdök was supported by The Scientific and Technological Research Council of Turkey 2219 post-doctoral research grant. Sequencing was performed at the National Genomics Infrastructure (NGI) Stockholm. Computational analyses were performed at the Uppsala Multidisciplinary Center for Advanced Computational Science (UPPMAX) under projects SNIC 2018/8-150 and SNIC 2018/8-248.

## Author contributions

M.A.M proposed the idea for the study. B.N. provided material and information on the Huseby Klev site. N.K. performed sampling and lab work, E.K. performed bioinformatic analyses, and they both contributed equally. H.D. conducted the lithic analysis. N.K., E.K., H.D., M.A.M. and P.P wrote the manuscript and supplements. P.P. and A.G. supervised the work.

---

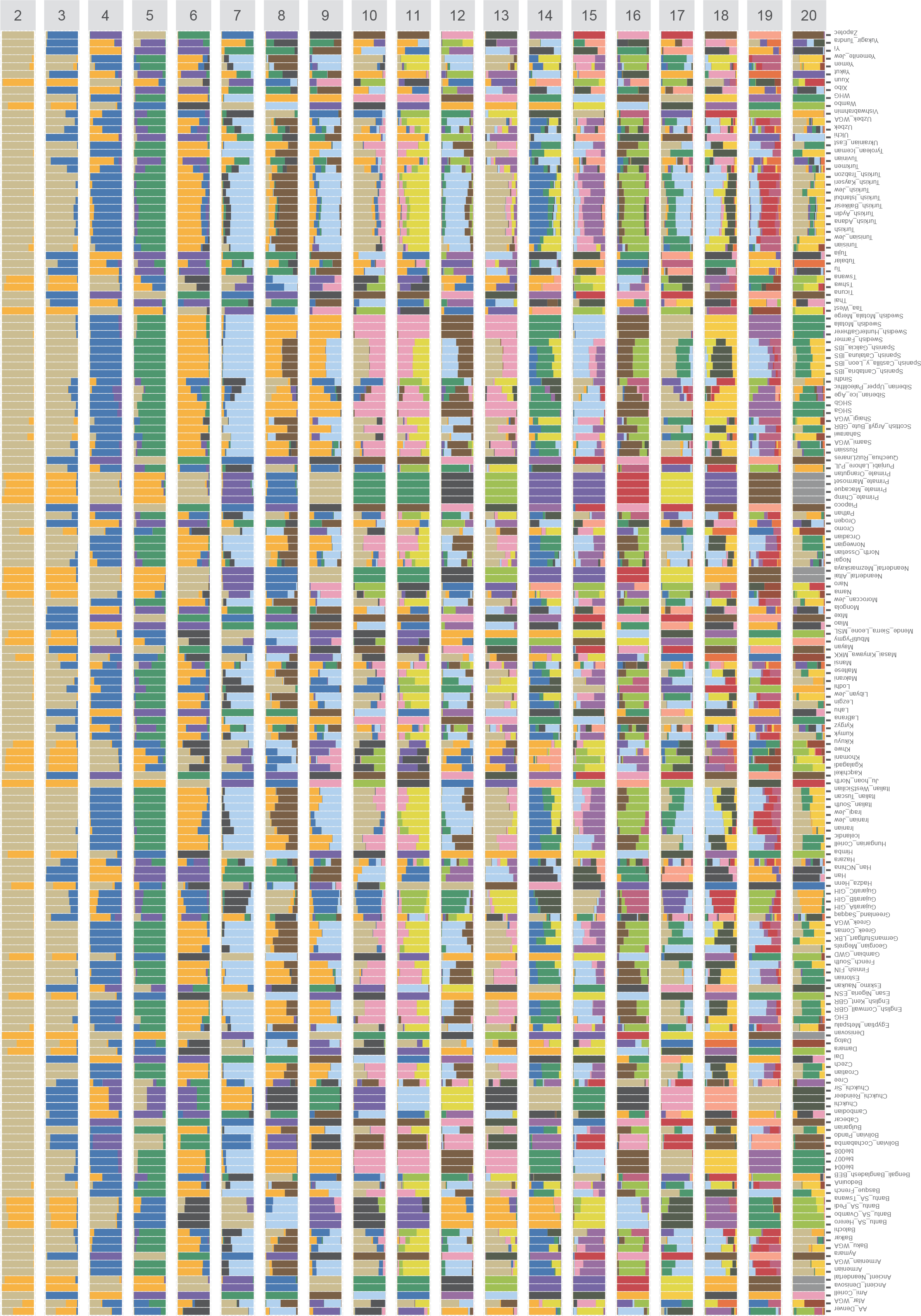

**Table S2.1.**
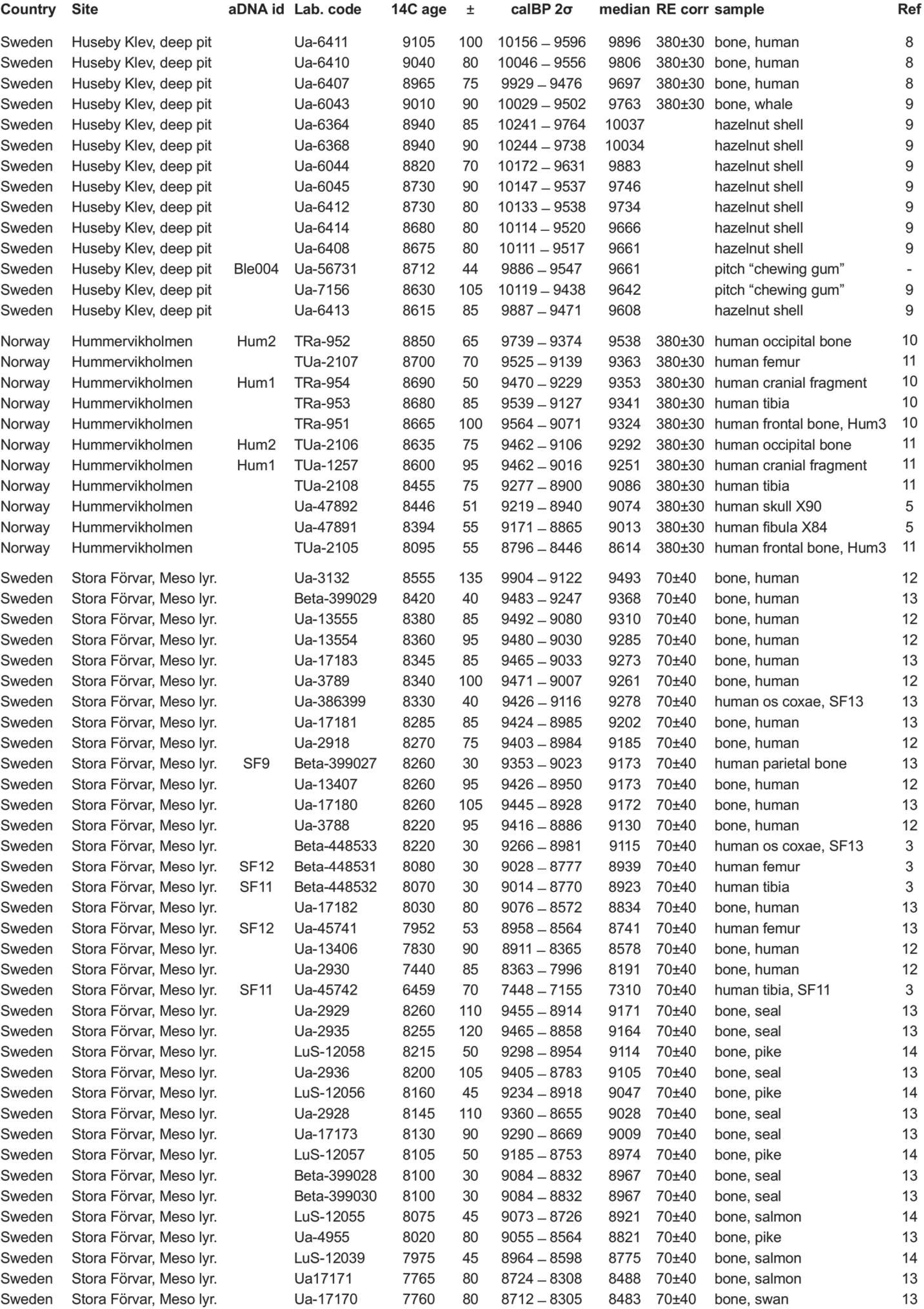
Radiocarbon dates from SHG contexts

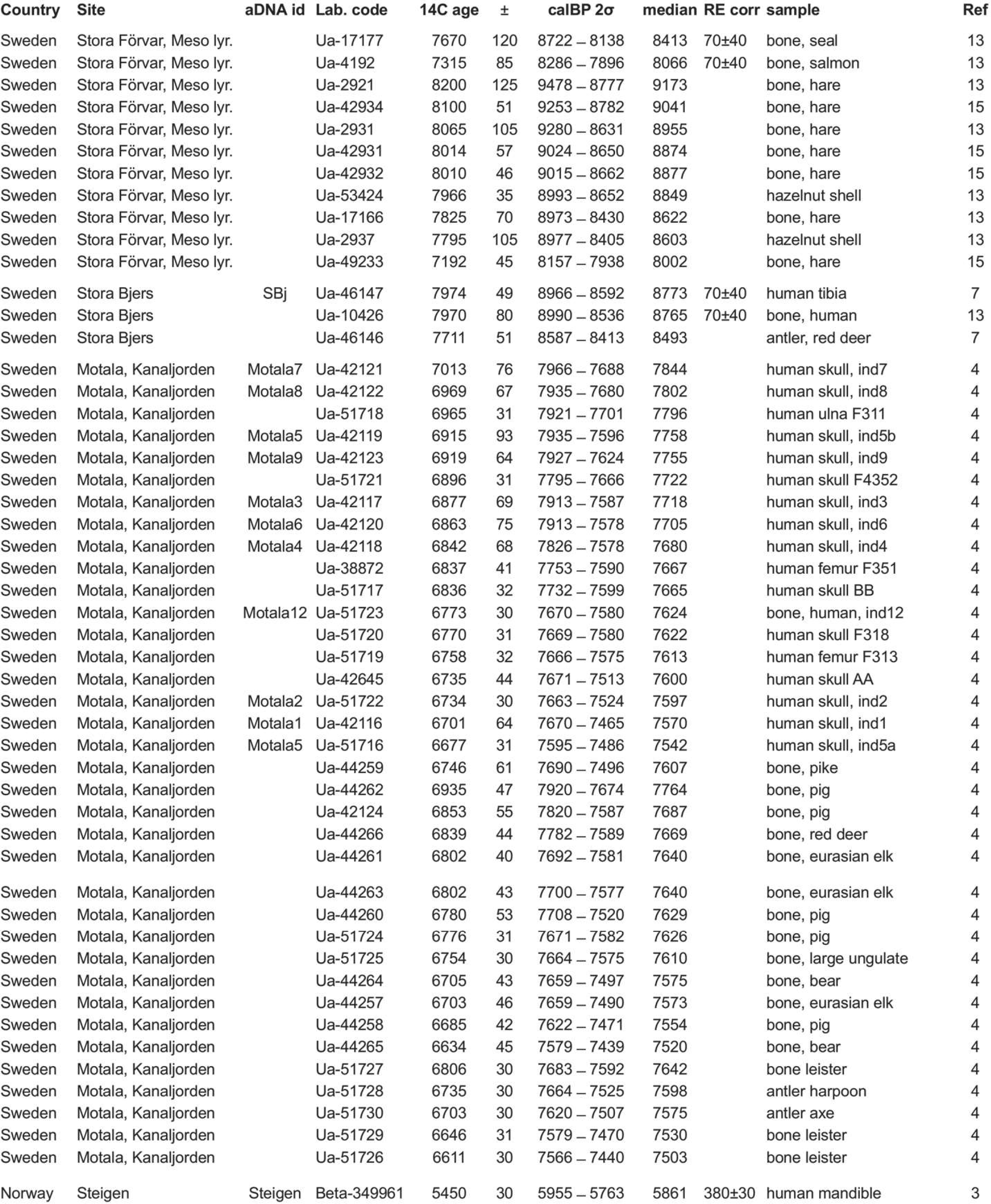

