## Supplementary material for "Ancient DNA from chewing gums connects material culture and genetics of Mesolithic hunter-gatherers in Scandinavia"

| Sequence id | ble004mergedStrict | | ble007 | | ble008mergedStrict | |
|---|---|---|---|---|---|---|
| **Final Haplogroup** | U5a2d | | U5a2d | | U5a2d | |
| **HaploFind prediction** | U5a2d | | U5a2d | | U5a2d | |
| **HaploFind score** | 1 | | 1 | | 1 | |
| **Haplogrep Prediction** | U5a2d | | U5a2d | | U5a2+16294 | |
| **Overall rank_ Haplogrep** | 0.9796 | | 0.8825 | | 1.1275 | |
| **Missing SNPs reported by Haplofind** | 195T, 2758G, 7146A, 7521G, 11467G, 12705C, 14793G | | 152T, 7146A, 7521G, 13506C, 13650C, 14793G, 16129G, 16270T | | 3197C, 11467G, 14793G, 16256T, 16270T | |
| | SNPs_reported_by_HaploFind | Polymorphisms (Haplogrep) | SNPs_reported_by_HaploFind | Polymorphisms (Haplogrep) | SNPs_reported_by_HaploFind | Polymorphisms (Haplogrep) |
| | 1_6del | 103R | 1_13del | 73G | 1_7del | 73R |
| | 146T | 128Y | 146T | 152C | 146T | 113Y |
| | 152T | 185R | 195T | 263G | 152T | 114Y |
| | 247G | 195Y | 247G | 386Y | 195T | 263G |
| | 311_315del | 228R | 303_309del | 433Y | 247G | 321N |
| | 769G | 263G | 311_312del | 458Y | 305_321del | 377N |
| | 825T | 462Y | 652del | 566Y | 377del | 387Y |
| | 1018G | 465Y | 660_661del | 709R | 769G | 413R |
| | 1083_1211del | 514Y | 769G | 750G | 825T | 447Y |
| | 1465_1481del | 541Y | 825T | 805Y | 1018G | 709R |
| | 1484_1485del | 544Y | 889del | 815Y | 1090_1097del | 750G |
| | 1725del | 658R | 920del | 983Y | 1224_1225del | 899R |
| | 2513del | 664R | 931_939del | 1375Y | 2758G | 1241Y |
| | 2885T | 750G | 943del | 1385Y | 2885T | 1278Y |
| | 3035del | 752Y | 1018G | 1497N | 3101_3113del | 1312Y |
| | 3106_3108del | 917Y | 1100_1225del | 1537Y | 3594C | 1412R |
| | 3197C | 1028R | 1233_1234del | 1838Y | 3896del | 1438G |
| | 3287del | 1303R | 1236del | 1845Y | 3928del | 1503R |
| | 3330del | 1375Y | 1447_1485del | 1888R | 4104A | 1576R |
| | 3333del | 1438G | 1493del | 1903N | 4312C | 1619Y |
| | 3594C | 1562R | 1497del | 2221Y | 4393del | 1714Y |
| | 3632del | 1579Y | 1574del | 2224Y | 4407_4408del | 1865Y |
| | 3983del | 1631Y | 1588_1603del | 2341Y | 4425_4812del | 1902Y |
| | 4104A | 1830R | 1627del | 2415Y | 5163_5218del | 2513Y |
| | 4159_4201del | 1845Y | 1901del | 2726Y | 5221del | 2496R |
| | 4312C | 2024Y | 1953_1988del | 3004Y | 5636_5680del | 2526Y |
| | 4374_4811del | 2136Y | 2501del | 3028R | 5961_6166del | 2706G |
| | 5061_5276del | 2209R | 2520del | 3032R | 6169del | 2774Y |
| | 5379del | 2272Y | 2758G | 3066Y | 6531del | 2821Y |
| | 5535_5778del | 2314Y | 2885T | 3083W | 7146A | 2920Y |
| | 5780_5781del | 2375Y | 2896del | 3149Y | 7256C | 3021M |
| | 5881del | 2706G | 2923del | 3197C | 7389del | 3009Y |
| | 5895_6145del | 2758R | 2927_2943del | 3204Y | 7392del | 3030R |
| | 6276del | 2761Y | 3106_3109del | 3283R | 7406_7490del | 3121Y |
| | 6671del | 2867Y | 3197C | 3284R | 7503del | 3152Y |
| | 6779_6880del | 3047R | 3340del | 3325Y | 7521G | 3197Y |
| | 7018_7167del | 3123R | 3594C | 3341N | 7556_7591del | 3208Y |
| | 7256C | 3197C | 3960_4039del | 3375Y | 7661del | 3449Y |
| | 7287del | 3245Y | 4104A | 3449Y | 7843G | 3570Y |
| | 7320_7782del | 3391R | 4125_4252del | 3506Y | 8320_8328del | 3797Y |
| | 7785_7789del | 3442Y | 4271del | 3712R | 8340del | 4228Y |
| | 7820del | 3445Y | 4312C | 3757Y | 8468C | 4252Y |
| | 7843G | 3567Y | 4350_4850del | 3867Y | 8655C | 4834Y |
| | 8273_8377del | 3573Y | 5044_5310del | 3904Y | 8701A | 4975R |
| | 8385del | 3589Y | 5368del | 4886Y | 8823_8950del | 5110Y |
| | 8468C | 3798Y | 5529_5777del | 4908Y | 8963del | 5434Y |
| | 8591_8623del | 4869Y | 5844_6199del | 4909Y | 9099del | 5462Y |
| | 8637del | 4972R | 6287_6305del | 4936Y | 9477A | 5481Y |
| | 8655C | 6222Y | 6308del | 5444Y | 9540T | 5864R |
| | 8701A | 6277N | 6402_6403del | 5467Y | 10398A | 5916Y |
| | 8767_8943del | 6291Y | 6600del | 6223Y | 10589del | 6311Y |
| | 8947del | 6368Y | 6602del | 6226Y | 10664C | 6325Y |
| | 8970del | 6557Y | 6769_6863del | 6349Y | 10688G | 6486Y |
| | 9112del | 7843G | 7000_7184del | 6370Y | 10810T | 6596Y |
| | 9126_9214del | 8032Y | 7256C | 6371Y | 10873T | 6669Y |
| | 9257del | 8117Y | 7388_7635del | 6623Y | 10915T | 6718R |
| | 9264_9265del | 8156R | 7719_7793del | 6686Y | 11914G | 6823Y |
| | 9477A | 8189R | 7843G | 6863N | 12159del | 6884Y |
| | 9540T | 8421Y | 7920_8085del | 7249Y | 12308G | 7028T |
| | 10197del | 8480Y | 8219del | 7843G | 12372A | 7147Y |
| | 10268del | 8513Y | 8293_8388del | 7853R | 12705C | 7185Y |
| | 10398A | 8684Y | 8397del | 7896R | 13105A | 7223Y |
| | 10664C | 9031Y | 8406del | 8150Y | 13276A | 7356R |
| | 10688G | 9074Y | 8468C | 8156R | 13506C | 7816Y |
| | 10810T | 9477A | 8578_8591del | 8464Y | 13617C | 7817Y |
| | 10873T | 9562Y | 8594del | 8513Y | 13650C | 7858Y |
| | 10915T | 9565R | 8605del | 8697R | 13964del | 7981Y |
| | 10968del | 10019Y | 8655C | 9042Y | 15172del | 8006Y |
| | 11430del | 10090Y | 8701A | 9281Y | 15268del | 8030Y |
| | 11914G | 10091Y | 8772_9010del | 9477A | 16129G | 8273Y |
| | 12019del | 10120Y | 9015del | 9477A | 16187C | 8398Y |
| | 12159_12175del | 10172R | 9118_9261del | 9746R | 16189T | 8406Y |
| | 12177del | 10247R | 9383_9459del | 9818Y | 16192T | 8514Y |
| | 12308G | 10268N | 9477A | 10314Y | 16223C | 8622Y |
| | 12372A | 10292Y | 9505del | 10463Y | 16230A | 8697R |
| | 12435del | 10293Y | 9540T | 10466Y | 16278C | 9011Y |
| | 13105A | 10466Y | 9592_9654del | 10751Y | 16311T | 9157R |
| | 13276A | 10769Y | 9693del | 10812Y | 16379del | 9289Y |

| | | | | | | |
|---|---|---|---|---|---|---|
| | 13477del | 10845Y | 9947del | 10938Y | 16407del | 9477A |
| | 13506C | 10944Y | 10090_10092del | 11054Y | 16519T | 9532Y |
| | 13617C | 11145Y | 10094del | 11174Y | 16526A | 9853Y |
| | 13650C | 11150R | 10198del | 11405Y | 16564_16569del | 10088Y |
| | 14565del | 11215Y | 10398A | 11467G | 16189T | 10091Y |
| | 16129G | 11227Y | 10534_10542del | 11515Y | 16192T | 10186Y |
| | 16187C | 11246R | 10664C | 11702Y | 16223C | 10268Y |
| | 16189T | 11279Y | 10688G | 11719A | 16230A | 10542Y |
| | 16192T | 11393Y | 10695del | 11858Y | 16278C | 10677R |
| | 16223C | 11467R | 10810T | 12109Y | 16311T | 10849Y |
| | 16230A | 11574Y | 10873T | 12123Y | 16519T | 10923Y |
| | 16256T | 11592R | 10915T | 12308G | 16526A | 10965Y |
| | 16270T | 11602Y | 11467G | 12372A | 16543del | 11063Y |
| | 16278C | 11675Y | 11489_11490del | 12485Y | 16564_16569del | 11217Y |
| | 16311T | 11719A | 11497_11498del | 12572R | | 11294Y |
| | 16519T | 11848Y | 11637_11659del | 12618R | | 11467R |
| | 16526A | 11994Y | 11914G | 12665Y | | 11515Y |
| | 16568_16569del | 12018N | 12308G | 12910Y | | 11537Y |
| | | 12025Y | 12372A | 13128Y | | 11539Y |
| | | 12276R | 12705C | 13328Y | | 11598Y |
| | | 12281Y | 12813del | 13351Y | | 11602Y |
| | | 12308G | 12852del | 13506Y | | 11719A |
| | | 12372A | 13052_13057del | 13617C | | 11813Y |
| | | 12514R | 13063del | 13662Y | | 11902R |
| | | 12548Y | 13105A | 13702Y | | 11942Y |
| | | 12705Y | 13199_13208del | 13704Y | | 12303Y |
| | | 12756R | 13211_13223del | 13828Y | | 12308G |
| | | 12869R | 13276A | 13908Y | | 12372A |
| | | 13035Y | 13617C | 13934Y | | 12411Y |
| | | 13128Y | 14051del | 14081R | | 12417Y |
| | | 13129Y | 14227_14228del | 14105Y | | 12741Y |
| | | 13204R | 14580_14601del | 14129Y | | 12858Y |
| | | 13325R | 15598del | 14263Y | | 12942Y |
| | | 13438Y | 16187C | 14266Y | | 12968Y |
| | | 13559Y | 16189T | 14410R | | 12969Y |
| | | 13583Y | 16192T | 14485Y | | 13026Y |
| | | 13617C | 16223C | 14671Y | | 13028Y |
| | | 13692Y | 16230A | 14751Y | | 13131Y |
| | | 13693Y | 16256T | 14766T | | 13368R |
| | | 13785Y | 16278C | 14793R | | 13617C |
| | | 13854Y | 16311T | 14905R | | 13625Y |
| | | 13948Y | 16519T | 15035Y | | 13694Y |
| | | 14003Y | 16526A | 15134R | | 13845Y |
| | | 14016R | 16562_16569del | 15179R | | 14335Y |
| | | 14055Y | | 15326G | | 14410R |
| | | 14124Y | | 15439Y | | 14529Y |
| | | 14142Y | | 15516Y | | 14620Y |
| | | 14252Y | | 15526Y | | 14766T |
| | | 14253Y | | 15661Y | | 14793R |
| | | 14340Y | | 15928R | | 14905R |
| | | 14343Y | | 16056Y | | 15037Y |
| | | 14427Y | | 16129R | | 15326G |
| | | 14448Y | | 16148Y | | 15334Y |
| | | 14519Y | | 16192T | | 15381Y |
| | | 14555Y | | 16256T | | 15433Y |
| | | 14565N | | 16270Y | | 15490Y |
| | | 14637Y | | 16294Y | | 15535Y |
| | | 14644Y | | 16295Y | | 15539Y |
| | | 14766T | | 16324Y | | 15619Y |
| | | 14793R | | 16388R | | 15739Y |
| | | 15016Y | | 16425Y | | 15928R |
| | | 15082Y | | 16526A | | 16036R |
| | | 15107Y | | | | 16052Y |
| | | 15218R | | | | 16192T |
| | | 15303Y | | | | 16256Y |
| | | 15326G | | | | 16270Y |
| | | 15375R | | | | 16294Y |
| | | 15490Y | | | | 16324Y |
| | | 15569Y | | | | 16457R |
| | | 15767Y | | | | 16526A |
| | | 15781Y | | | | 15037Y |
| | | 15815Y | | | | 15326G |
| | | 15939Y | | | | 15304N |
| | | 16192T | | | | 15286N |
| | | 16256T | | | | 15334Y |
| | | 16270T | | | | 15381Y |
| | | 16399R | | | | 15409Y |
| | | 16526A | | | | 15433Y |
| | | | | | | 15434Y |
| | | | | | | 15468Y |
| | | | | | | 15535Y |
| | | | | | | 15516Y |
| | | | | | | 15539Y |
| | | | | | | 15619Y |
| | | | | | | 15632Y |
| | | | | | | 15664Y |
| | | | | | | 15753Y |
| | | | | | | 15928R |
| | | | | | | 16036R |

| | | | | | | |
|---|---|---|---|---|---|---|
| | | | | | | 16052Y |
| | | | | | | 16147Y |
| | | | | | | 16192T |
| | | | | | | 16256Y |
| | | | | | | 16270Y |
| | | | | | | 16294Y |
| | | | | | | 16324Y |
| | | | | | | 16376Y |
| | | | | | | 16408Y |
| | | | | | | 16526A |
| | | | | | | 16543N |
