## Supplementary material for "Ancient DNA from chewing gums connects material culture and genetics of Mesolithic hunter-gatherers in Scandinavia"

F4

| PopA | PopB | PopX | PopY | F4 | SE | Z-score | nSites | nBlocks | nCHRa | nCHRB | nCHRX | nCHRy |
|---|---|---|---|---|---|---|---|---|---|---|---|---|
| Primate_Chim | ble004 | EHG | WHG | 0.01771148 | 0.02213388 | 0.80019772 | 2239 | 430 | 2 | 2 | 6 | 6 |
| Primate_Chim | ble007 | EHG | WHG | 0.00352113 | 0.02315508 | 0.15206712 | 2090 | 438 | 2 | 2 | 6 | 6 |
| Primate_Chim | ble008 | EHG | WHG | 0.02196558 | 0.01219181 | 1.80166641 | 7667 | 508 | 2 | 2 | 6 | 6 |
| Primate_Chim | ble004 | EHG | SHG | 0.02647897 | 0.01975763 | 1.34018937 | 2774 | 448 | 2 | 2 | 6 | 26 |
| Primate_Chim | ble007 | EHG | SHG | 0.02087439 | 0.02002955 | 1.04217997 | 2625 | 455 | 2 | 2 | 6 | 26 |
| Primate_Chim | ble008 | EHG | SHG | 0.03345158 | 0.01133265 | 2.95178646 | 9456 | 510 | 2 | 2 | 6 | 26 |
| Primate_Chim | ble004 | WHG | SHG | 0.01092498 | 0.01449675 | 0.75361618 | 3002 | 458 | 2 | 2 | 6 | 26 |
| Primate_Chim | ble007 | WHG | SHG | 0.01600133 | 0.01576499 | 1.01499157 | 2861 | 457 | 2 | 2 | 6 | 26 |
| Primate_Chim | ble008 | WHG | SHG | 0.01436639 | 0.00875846 | 1.64028648 | 10257 | 518 | 2 | 2 | 6 | 26 |
| Primate_Chim | ble004 | EHG | SHGb | 0.02764573 | 0.02109793 | 1.31035242 | 2585 | 444 | 2 | 2 | 6 | 20 |
| Primate_Chim | ble007 | EHG | SHGb | 0.02620157 | 0.02025954 | 1.29329517 | 2434 | 447 | 2 | 2 | 6 | 20 |
| Primate_Chim | ble008 | EHG | SHGb | 0.03551412 | 0.01141888 | 3.11012363 | 8821 | 508 | 2 | 2 | 6 | 20 |
| Primate_Chim | ble004 | EHG | SHGa | 0.02162952 | 0.02333939 | 0.92673867 | 1979 | 422 | 2 | 2 | 6 | 6 |
| Primate_Chim | ble007 | EHG | SHGa | 0.00998336 | 0.02393934 | 0.41702736 | 1851 | 419 | 2 | 2 | 6 | 6 |
| Primate_Chim | ble008 | EHG | SHGa | 0.03084046 | 0.01346439 | 2.29052108 | 6689 | 503 | 2 | 2 | 6 | 6 |
| Primate_Chim | ble004 | WHG | SHGb | 0.01315301 | 0.01584189 | 0.83026747 | 2844 | 455 | 2 | 2 | 6 | 20 |
| Primate_Chim | ble007 | WHG | SHGb | 0.02328217 | 0.01693177 | 1.37505791 | 2689 | 454 | 2 | 2 | 6 | 20 |
| Primate_Chim | ble008 | WHG | SHGb | 0.01508239 | 0.00911723 | 1.65427249 | 9694 | 514 | 2 | 2 | 6 | 20 |
| Primate_Chim | ble004 | WHG | SHGa | -0.00199203 | 0.01868726 | -0.1065984 | 2436 | 445 | 2 | 2 | 6 | 6 |
| Primate_Chim | ble007 | WHG | SHGa | 0.0019495 | 0.02019293 | 0.09654362 | 2302 | 437 | 2 | 2 | 6 | 6 |
| Primate_Chim | ble008 | WHG | SHGa | 0.01119726 | 0.01151763 | 0.97218397 | 8243 | 511 | 2 | 2 | 6 | 6 |
| Primate_Chim | ble004 | SHGb | SHGa | -0.01109329 | 0.01802357 | -0.61548785 | 2753 | 451 | 2 | 2 | 20 | 6 |
| Primate_Chim | ble007 | SHGb | SHGa | -0.01860966 | 0.0182369 | -1.02044007 | 2617 | 450 | 2 | 2 | 20 | 6 |
| Primate_Chim | ble008 | SHGb | SHGa | -0.002611 | 0.00979202 | -0.26664615 | 9350 | 517 | 2 | 2 | 20 | 6 |

| PopA | PopB | PopX | PopY | D | SE | Z-score | nSites | nBlocks | nCHRa | nCHRB | nCHRX | nCHRy |
|---|---|---|---|---|---|---|---|---|---|---|---|---|
| Primate_Chim | ble004 | EHG | WHG | 0.01771148 | 0.02213388 | 0.80019772 | 2239 | 430 | 2 | 2 | 6 | 6 |
| Primate_Chim | ble007 | EHG | WHG | 0.00352113 | 0.02315508 | 0.15206712 | 2090 | 438 | 2 | 2 | 6 | 6 |
| Primate_Chim | ble008 | EHG | WHG | 0.02196558 | 0.01219181 | 1.80166641 | 7667 | 508 | 2 | 2 | 6 | 6 |
| Primate_Chim | ble004 | EHG | SHG | 0.02647897 | 0.01975763 | 1.34018937 | 2774 | 448 | 2 | 2 | 6 | 26 |
| Primate_Chim | ble007 | EHG | SHG | 0.02087439 | 0.02002955 | 1.04217997 | 2625 | 455 | 2 | 2 | 6 | 26 |
| Primate_Chim | ble008 | EHG | SHG | 0.03345158 | 0.01133265 | 2.95178646 | 9456 | 510 | 2 | 2 | 6 | 26 |
| Primate_Chim | ble004 | WHG | SHG | 0.01092498 | 0.01449675 | 0.75361618 | 3002 | 458 | 2 | 2 | 6 | 26 |
| Primate_Chim | ble007 | WHG | SHG | 0.01600133 | 0.01576499 | 1.01499157 | 2861 | 457 | 2 | 2 | 6 | 26 |
| Primate_Chim | ble008 | WHG | SHG | 0.01436639 | 0.00875846 | 1.64028648 | 10257 | 518 | 2 | 2 | 6 | 26 |
| Primate_Chim | ble004 | EHG | SHGb | 0.02764573 | 0.02109793 | 1.31035242 | 2585 | 444 | 2 | 2 | 6 | 20 |
| Primate_Chim | ble007 | EHG | SHGb | 0.02620157 | 0.02025954 | 1.29329517 | 2434 | 447 | 2 | 2 | 6 | 20 |
| Primate_Chim | ble008 | EHG | SHGb | 0.03551412 | 0.01141888 | 3.11012363 | 8821 | 508 | 2 | 2 | 6 | 20 |
| Primate_Chim | ble004 | EHG | SHGa | 0.02162952 | 0.02333939 | 0.92673867 | 1979 | 422 | 2 | 2 | 6 | 6 |
| Primate_Chim | ble007 | EHG | SHGa | 0.00998336 | 0.02393934 | 0.41702736 | 1851 | 419 | 2 | 2 | 6 | 6 |
| Primate_Chim | ble008 | EHG | SHGa | 0.03084046 | 0.01346439 | 2.29052108 | 6689 | 503 | 2 | 2 | 6 | 6 |
| Primate_Chim | ble004 | WHG | SHGb | 0.01315301 | 0.01584189 | 0.83026747 | 2844 | 455 | 2 | 2 | 6 | 20 |
| Primate_Chim | ble007 | WHG | SHGb | 0.02328217 | 0.01693177 | 1.37505791 | 2689 | 454 | 2 | 2 | 6 | 20 |
| Primate_Chim | ble008 | WHG | SHGb | 0.01508239 | 0.00911723 | 1.65427249 | 9694 | 514 | 2 | 2 | 6 | 20 |
| Primate_Chim | ble004 | WHG | SHGa | -0.00199203 | 0.01868726 | -0.1065984 | 2436 | 445 | 2 | 2 | 6 | 6 |
| Primate_Chim | ble007 | WHG | SHGa | 0.0019495 | 0.02019293 | 0.09654362 | 2302 | 437 | 2 | 2 | 6 | 6 |
| Primate_Chim | ble008 | WHG | SHGa | 0.01119726 | 0.01151763 | 0.97218397 | 8243 | 511 | 2 | 2 | 6 | 6 |
| Primate_Chim | ble004 | SHGb | SHGa | -0.01109329 | 0.01802357 | -0.61548785 | 2753 | 451 | 2 | 2 | 20 | 6 |
| Primate_Chim | ble007 | SHGb | SHGa | -0.01860966 | 0.0182369 | -1.02044007 | 2617 | 450 | 2 | 2 | 20 | 6 |
| Primate_Chim | ble008 | SHGb | SHGa | -0.002611 | 0.00979202 | -0.26664615 | 9350 | 517 | 2 | 2 | 20 | 6 |
